# An ensemble method for predicting and designing of druggable proteins

**DOI:** 10.1101/2024.10.17.618792

**Authors:** Shipra Jain, Srijanee Gupta, Gajendra P. S. Raghava

**Affiliations:** Department of Computational Biology, Indraprastha Institute of Information Technology, Okhla Phase 3, New Delhi-110020, India

**Keywords:** Druggable Proteins, Therapeutics, Machine learning techniques, motifs

## Abstract

In the past, numerous proteins/peptides have been discovered, which have a wide range of therapeutic properties like anticancer, antimicrobial, antihypertensive. Only few hundreds of proteins are druggable (approved by US FDA), most of the proteins fails in clinical trials. In this study, an attempt had been made to understand properties of FDA approved proteins to develop models for predicting druggable proteins. Our main dataset 356 FDA approved proteins as positive dataset and equal number of randomly selected proteins as negative dataset. We used 80% data for training and 20% for independent validation, no protein in validation dataset have more than 40% similarity with any protein in training dataset. We deployed machine learning based models using a five-fold cross-validation and test on validation dataset. Our random forest-based model developed using SVC-L1 selected features obtained maximum performance AUC of 0.80 with MCC 0.61 on validation data. In addition to this, we performed MERCI-based motif analysis to find motifs in druggable proteins. Finally, we developed an ensemble-based method combining best performing machine learning model with motifs and achieved AUC 0.92 with MCC 0.83 on independent validation dataset. We developed a web server and standalone package ThPPred to facilitate scientific community in predicting and designing druggable proteins (https://webs.iiitd.edu.in/raghava/thppred/).

**Highlights:**

1. Analysis of FDA approved or druggable proteins
2. Discrimination of druggable and non-druggable proteins
3. Machine learning based models for predicting druggable molecules
4. Identification of motifs in druggable proteins
5. A web server for providing service to community

## Introduction

In the last three decades, there is an exponential growth in protein associated databases due to rapid advancement in next generation sequencing technologies. In the past, number of computational resources have been developed to facilitate scientific community in annotating protein’s structure, function and therapeutic properties^1,2^. Recently, field of therapeutic proteins/peptides have been emerged, which is evident from number of proteins approved by US-FDA for market use^3–5^. Proteins play a vital role in human physiology such as body growth factors, hormones, immune regulation, transmitters and metabolism^6^. It has many advantages over small molecules-based drugs that includes higher specificity for target, lower toxicity, environment friendly^7,8^. Therapeutic proteins are used to treat a wide range of diseases including several type of cancer, autoimmune diseases, diabetes mellitus, genetic abnormalities and hormone imbalances^4,9–13^.

Several computational methods to predict various therapeutic protein types have been presented in the past such as PTPD^14^, PrMFTP^15^, TPpred-ATMV^16^, TPpred-LE^17^, PreTP-2L^18^, Pep-CNN^19^, PreTP-Stack^20^, *q*-FP^21^, TP-MV^22^, PPTPP^23^, PEPred-Suite^24^, PreTP-EL^25^ etc. However, these models have concentrated on proteins/peptides specific properties such as anti-angiogenic peptides, anti-bacterial peptides, anti-cancer peptides, anti-inflammatory peptides, anti-viral peptides, cell-penetrating peptides, polystyrene surface binding peptides, anti-parasitic peptides and quorum sensing peptides rather than proteins/peptides that have been recognized and approved by FDA as drug molecules. Thus, creating a pressing need for developing a prediction model with reported therapeutic proteins/peptides that gives better insights in identifying potential therapeutic candidates for future drug discoveries. In order to reduce the cost of development of drugs, prediction models are proven to be highly imperative.

With these facts in mind, a systematic attempt has been made in this study to create an in-silico technique for predicting therapeutic proteins and peptides utilizing experimentally validated therapeutic proteins/peptides for identification and classification of proteins as therapeutic and non-therapeutic. We believe that ThPPred is a right initiative in this direction, and would be highly useful for the scientific community working in the drug discovery domain, leading to fast and accurate screening of the therapeutic proteins/peptides. The web server and standalone version of the ThPPred is available at https://webs.iiitd.edu.in/raghava/thppred/.

## Materials and Methods

### Compilation of datasets

#### Therapeutic proteins as positive dataset

In this study, we downloaded 381 FDA approved therapeutic proteins/peptides data reported in ThpDB2^2^. We removed sequences having less than 30 amino acids or more than 1500 amino acids. Finally, we have 356 sequences which have length between 30-1500 and having no non-natural amino acids, which were referred to as a positive dataset in this paper.

#### Non-Therapeutic proteins as negative dataset

We searched protein in from Swiss-Prot with “reviewed” which have therapeutic properties using following keywords anti-antiogenic, anti-bacterial, anti-cancer, anti-inflammatory, anti-viral, cell-penetrating, polystyrene surface binding, and quorum sensing etc. After removing duplicate entries, we were left with 83655 sequences. Of these proteins/peptides, sequences of length 30-1500 were filtered, and we’re left with 83313 sequences. We got non-redundant sequences after removing redundant proteins from these proteins using CD-HIT software with 40% cut-off^26–29^. We randomly selected 356 non-redundant sequences to create our main dataset. Our main dataset contains 356 therapeutic and 356 non-redundant proteins. Similarly, we randomly 3560 non-redundant proteins to create realistic dataset. Our realistic dataset contains 356 therapeutic and 3560 non-therapeutic proteins.

### Non-Redundant Dataset Stratification

Ideally a dataset should have non-redundant proteins where no two proteins have more than 40% sequence similarity with each other. Though our non-therapeutic proteins are non-redundant, but our therapeutic proteins have redundancy. Number of therapeutic proteins reduced drastically if we remove similar proteins. We applied clustering technique to reduce redundancy between training and validation dataset without reducing number of therapeutic proteins. Similar technique has been used in previous studies to solve this problem of redundancy^27,28^. In this study, we implied a similar approach to bifurcate our data in 80% training and 20% independent validation dataset, where no protein in training data had more than 40% similarity with any sequence in the validation dataset. In order to achieve this, we created clusters using CD-HIT software at a 40% sequence similarity for both datasets^30^. Using this technique, we obtained a total of 133 clusters in the positive dataset and 139 clusters in negative data of the main dataset. These clusters were divided in such a way that 80% data goes to training dataset and 20% data to validation set. Training data was further divided into 5 sets where each set have 20% data of training dataset. As division of data was at cluster level so no protein in training dataset have similarity with proteins in validation dataset. The graphical representation of steps followed for segregating non redundant dataset is depicted in Figure 2.

**Figure 1:**
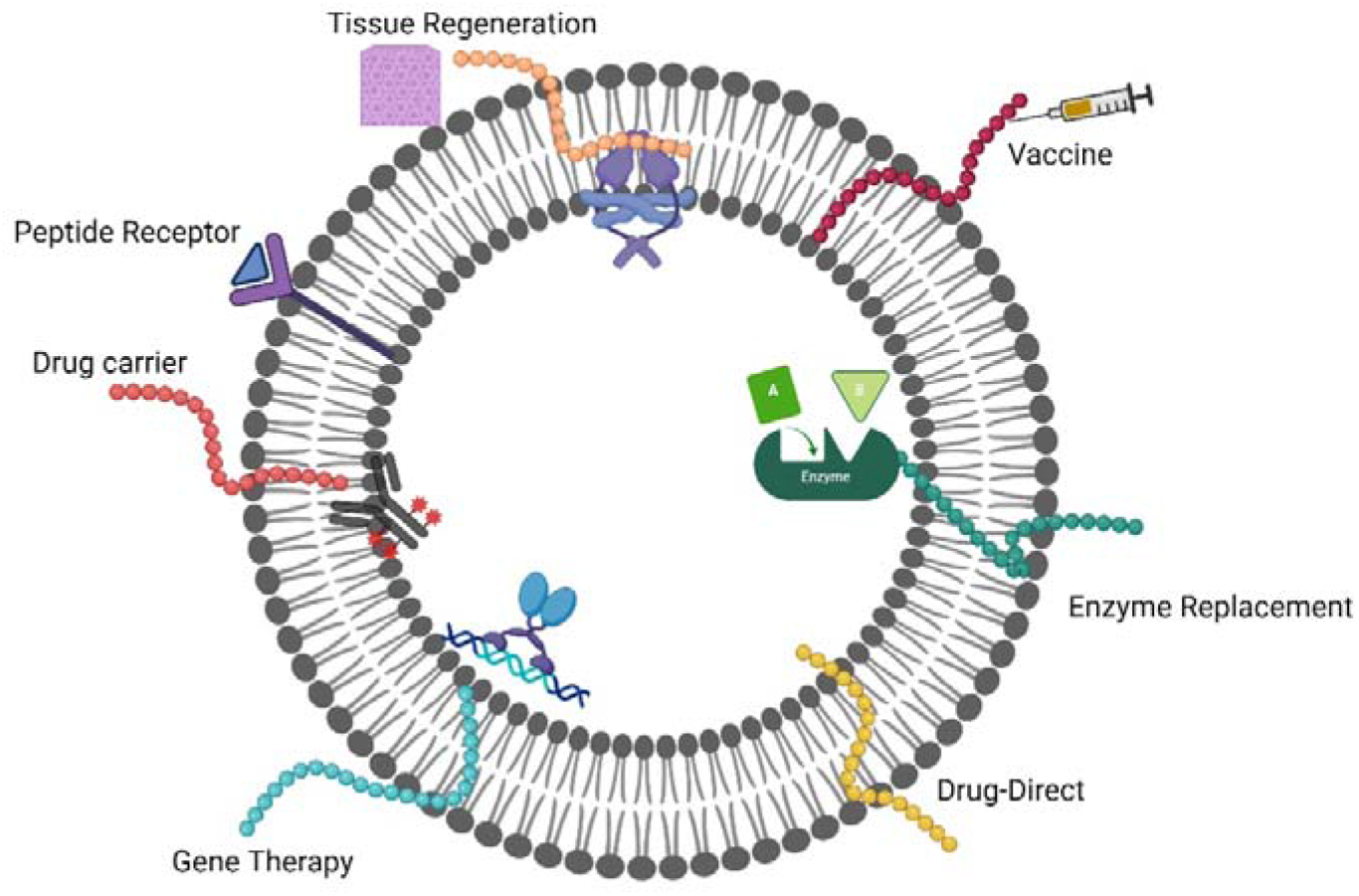
Figure depicting applications of therapeutic proteins/peptides.

**Figure 2:**
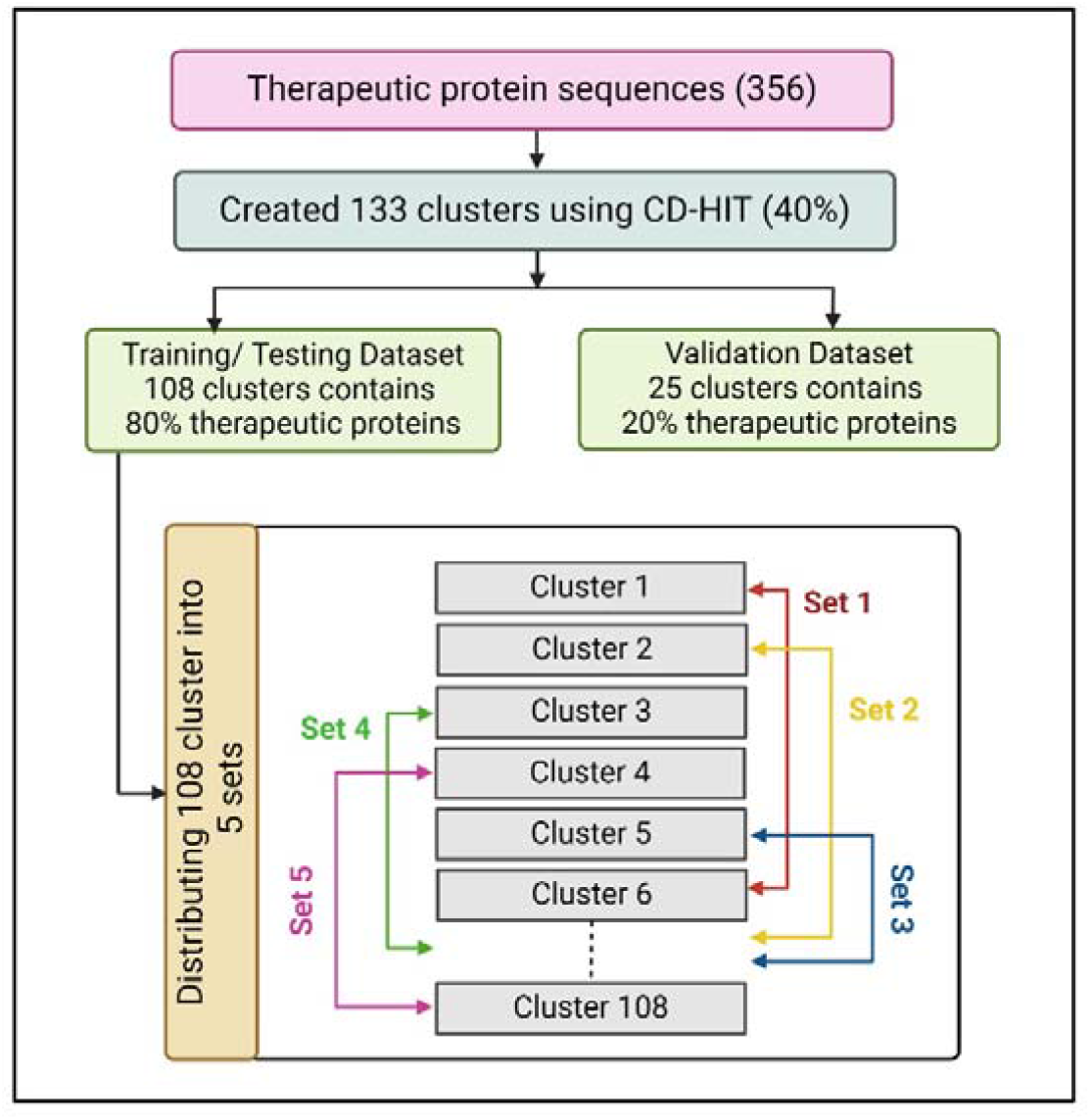
Flowchart shows division of clusters into training and validation dataset.

### Feature generation & selection

In this study, we used Pfeature for generating a wide range features (around 9189 features) corresponding to a protein^31^. As all features are not relevant thus, we used following feature selection techniques to select relevant features.

1. **SVC-L1:** We used Scikit-learn package for selecting features using support vector classifier with linear kernel and L1 regularization.
2. **Variance Threshold:** A feature’s variance is a measurement of how far it deviates from its mean value. A feature with low variance is one whose values are closely grouped around the mean. We selected only those features which have variance more than a threshold.
3. **Correlation Coefficient:** We removed all those features which are highly correlated with each other as all provide similar information.
4. **PCA:** In addition to feature selection, we also used feature merging technique principal component analysis (PCA). In these techniques, we merge all feature and selected top principal components as features.

### Cross-validation and performance metrics

In this study, data was divided into an 80:20 ratio, where 80% make up the training datasets and 20% serve as the validation datasets. Five-fold cross-validation while building our model on the 80% of training dataset was opted in this study. The 80% training data is divided into five folds for internal validation, with four folds utilized for training and the last fold for testing. Five rounds of the same process are iterated in order to test each of the five folds at least once. The average of the common evaluation metrics including Sensitivity, Specificity, Accuracy, and Matthews Correlation Coefficient (MCC) as threshold-dependent metrics, and the area under the receiver operating characteristic curve (AUC) is a parameter that is independent of threshold. Models were then evaluated based on the predicted performance over the unseen validation set. This standard procedure has been successfully implemented in several past machine learning (ML)-based research^27,29,32–35^.

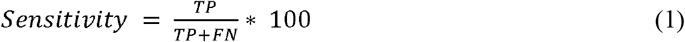

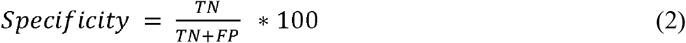

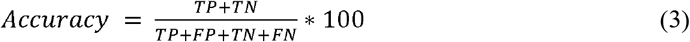

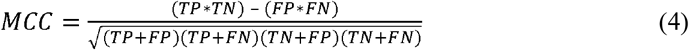

Where, FP is false positive, FN is false negative, TP is true positive and TN is true negative.

### Machine Learning based classifiers

In this study, machine learning techniques have been implemented for developing classification model using python package scikit-learn. We developed models using following techniques Random Forest (RF)^36^, Decision Tree^37^, Logistic Regression (LR)^38^, XGBoost (XGB)^39^, k-nearest neighbours (KNNs)^40^, and Support Vector Classifier^41^. Hyperparameters of these classifiers were optimized to obtain best prediction accuracy. The illustrative workflow of ThpPred is depicted in Figure 3.

**Figure 3:**
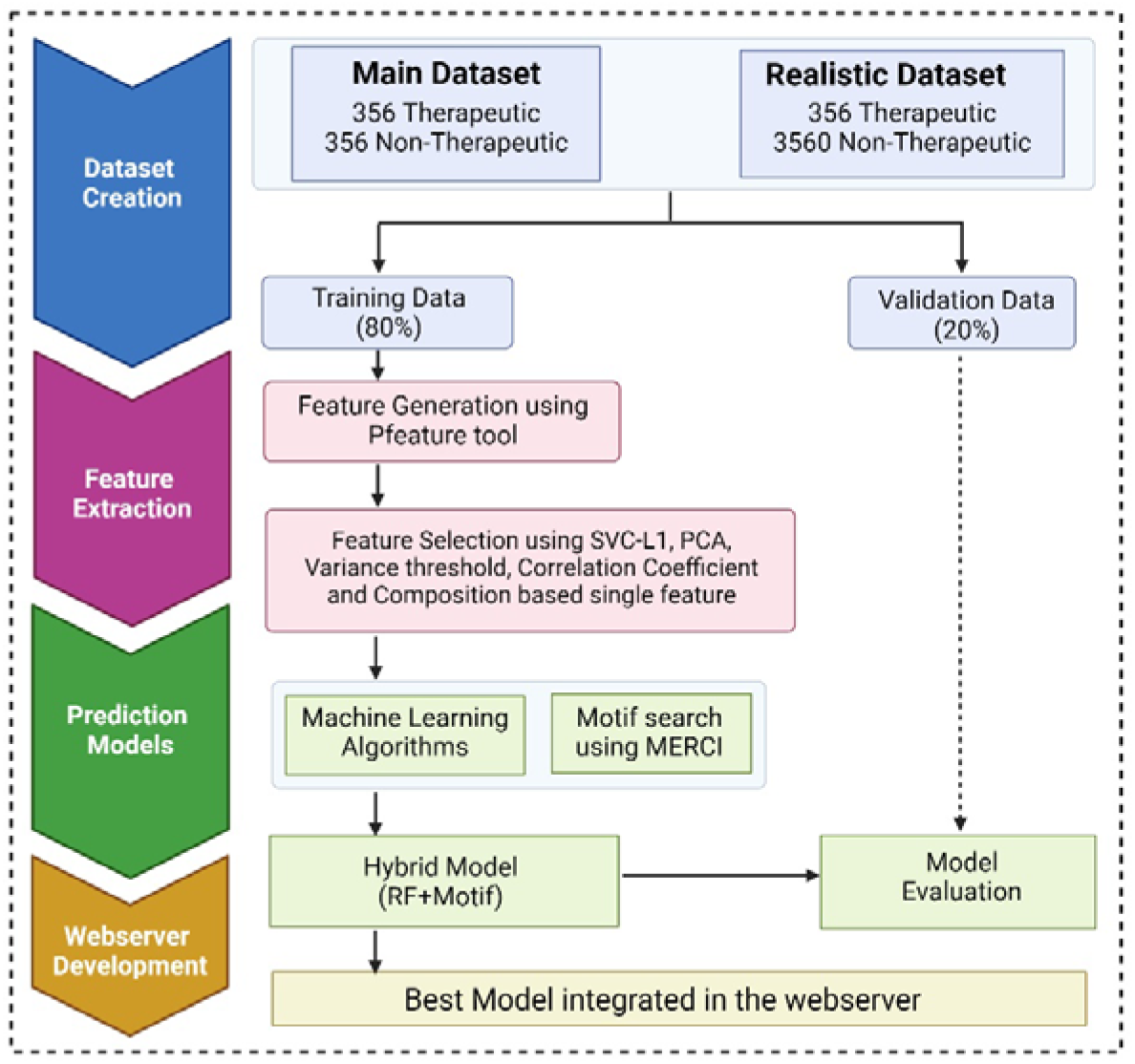
Pictorial representation of ThpPred workflow.

### Motif Scanning

We implement software Motif-EmeRging and the Classes-Identification (MERCI) for discovering motifs or abundant patterns in therapeutic proteins. This is a Perl based tool heavily used in number of previous studies for detecting motifs^42–44^.

### Hybrid Approach

In this study, we have also implemented an ensemble approach for classifying therapeutic and non-therapeutic proteins/peptides. In this approach, we developed a hybrid method of combining the best performing machine learning model with a higher coverage and motif-based analysis done using the MERCI tool with higher precision prediction capabilities. First of all, we assigned ‘+0.5’ weight for the positive predictions (therapeutic proteins), ‘−0.5’ for negative predictions (non-therapeutic proteins) and ‘0’ for no hits. Secondly, the same protein sequence was classified using the MERCI tool. We assigned the score of ‘+0.5’ if the motifs were found and ‘0’ if the motifs were not found. The overall score was calculated at different threshold values, and the protein/peptide sequence is predicted as therapeutic and non-therapeutic. This hybrid approach has been extensively deployed in various past studies^27,29^.

## Results

### Compositional Analysis

We perform compositional analysis of therapeutic proteins to understand their insights. As depicted in Figure 4, we observed that in the main dataset composition of Cysteine, Lysine, Asparagine, Serine, Valine and Tyrosine are higher in therapeutic proteins as compared to non-therapeutic proteins/peptides. Whereas, Alanine, Isoleucine and Arginine are prominent in non-therapeutic proteins. On top of that, similar trend has been observed for the realistic dataset.

**Figure 4:**
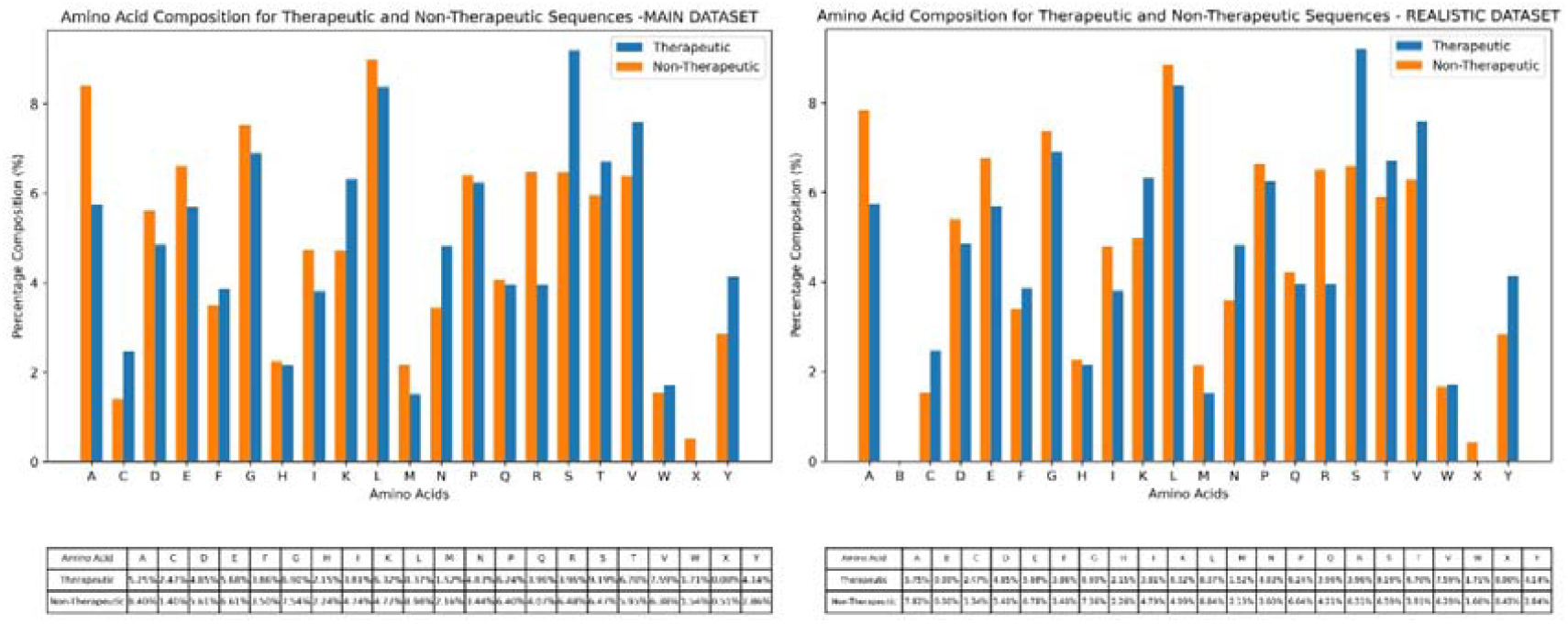
Comparison of amino acid composition of therapeutic and non-therapeutic proteins in main and realistic dataset.

### Machine learning based models

In this study, relevant features were extracted using several feature selection techniques such as SVC-L1, Variance Threshold, Principal Component Analysis, correlation coefficient and composition-based feature such as AAC, DPC & TPC. The complete list of features extracted using various methods can be referred in Supplementary Table S1. Various machine learning models were developed using scikit-learn python library. The performance of machine learning based models developed using selected 60 features obtained from SVC-L1, is shown in Table 1. As shown in Table 1, these models have huge variation from AUC 0.58 to 0.80. Our best model based on Random Forest perform better than other models and achieved maximum AUC 0.80 with MCC 0.61 and Accuracy 0.80 on a validation dataset. Detailed list of the results for various classifiers can be referred to Supplementary Table S2.

**Table 1:**
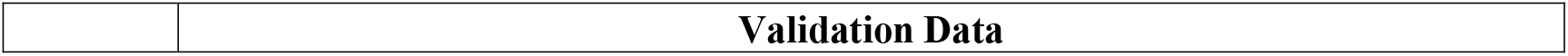
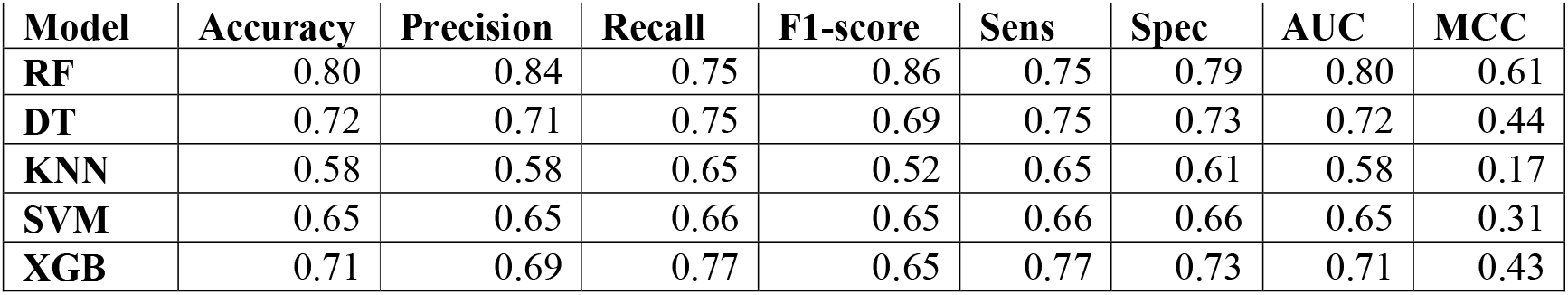
The performance of machine learning based models on validation or independent dataset, models developed using SVC-L1 selected features.

| Model | Accuracy | Precision | Recall | F1-score | Sens | Spec | AUC | MCC |
| --- | --- | --- | --- | --- | --- | --- | --- | --- |
| RF | 0.80 | 0.84 | 0.75 | 0.86 | 0.75 | 0.79 | 0.80 | 0.61 |
| DT | 0.72 | 0.71 | 0.75 | 0.69 | 0.75 | 0.73 | 0.72 | 0.44 |
| KNN | 0.58 | 0.58 | 0.65 | 0.52 | 0.65 | 0.61 | 0.58 | 0.17 |
| SVM | 0.65 | 0.65 | 0.66 | 0.65 | 0.66 | 0.66 | 0.65 | 0.31 |
| XGB | 0.71 | 0.69 | 0.77 | 0.65 | 0.77 | 0.73 | 0.71 | 0.43 |

### Motif Analysis

In addition to the machine learning classifiers, we have incorporated motif analysis over our data using MERCI software. Using this tool, we identified 25 exclusive motifs present in therapeutic proteins/ peptides and 20 exclusive motifs in non-therapeutic proteins/ peptides for main dataset. Top 5 motifs occurring in positive and negative sequences of the main dataset are listed in Table 2, and a detailed list of all selected motifs for both datasets can be referred to in Supplementary Table S4.

**Table 2:** List of top five motifs obtained using MERCI software in therapeutic and non-therapeutic proteins in main dataset.

| Motifs exclusively present in Therapeutic Proteins |  | Motifs exclusively present in non-therapeutic proteins |  |
| --- | --- | --- | --- |
| Motifs | Occurrence | Motifs | Occurrence |
| sulfurbasicgapSneutral | 135 | PVchargedP | 135 |
| YaromaticC | 133 | PVchargedPsurface | 132 |
| NeutralKNneutral | 133 | PVchargedPpolar | 121 |
| WgapQG | 132 | PVacidicP | 108 |
| Qaliphaticaromaticneutralaliphatic | 132 | PVchargedPneutral | 106 |

### Performance evaluation using Ensemble method

In order to capture molecular insights of our therapeutic peptides data, we developed an ensemble method combining our best performing machine learning model with higher coverage and the motif-based analysis done using the MERCI tool with higher precision prediction. Using this strategy, we attempted to develop a hybrid prediction method with higher accuracy combining capabilities of both machine learning as well as motif analysis. We achieved highest 0.92 AUC score for Random Forest classifier developed over main data using SVC-L1 feature extraction technique. Performance achieved using various classifiers over main dataset are reported in Table 3.

**Table 3:** Performance evaluation for ensemble method combining SVC-L1 feature selection technique-based ML models with motif analysis for the main dataset.

| Model | Validation Data |  |  |  |  |  |  |  |
| --- | --- | --- | --- | --- | --- | --- | --- | --- |
|  | Accuracy | Precision | Recall | F1-score | Sens | Spec | AUC | MCC |
| **RF** | 0.92 | 0.92 | 0.92 | 0.92 | 0.92 | 0.92 | 0.92 | 0.83 |
| **DT** | 0.89 | 0.86 | 0.94 | 0.85 | 0.94 | 0.90 | 0.89 | 0.79 |
| **KNN** | 0.85 | 0.82 | 0.90 | 0.80 | 0.90 | 0.86 | 0.85 | 0.71 |
| **SVM** | 0.86 | 0.86 | 0.86 | 0.86 | 0.86 | 0.86 | 0.86 | 0.72 |
| **XGB** | 0.91 | 0.90 | 0.92 | 0.90 | 0.92 | 0.91 | 0.91 | 0.82 |

### Prediction classifier performance over realistic dataset

In this study, we have applied similar approach as main dataset for developing machine learning models over ten times the negative dataset as compared to positive data tagged as a realistic dataset. For realistic dataset, Support Vector based classifier build over 91 relevant features selected using SVC-L1 technique outperformed other classifiers. The maximum AUC reported over independent validation was 0.78. Performance achieved using other classifiers are reported in Supplementary Table S3. In addition to machine learning based classifiers, we have developed ensemble method similar to main dataset over realistic data. Using this approach, AUC score for Support Vector classifier using SVC-L1 feature extraction technique was reported to be same as 0.78. Detailed results for all classifiers implemented over realistic dataset can be referred in Supplementary Table S5.

### Design and implementation of a web server

In order to serve the scientific community, we have provided a user-friendly web server, which can be accessed over https://webs.iiitd.edu.in/raghava/thppred/. Best performing machine learning models proposed in this study have been incorporated in the webserver. It gives users flexibility to choose machine learning models trained on either main or realistic dataset, desired threshold value and upload FASTA sequence file in one go. The web server integrates the key components, including (i) prediction, (ii) motif scan, and (iii) therapeutic protein design module. The ‘prediction module’ enables users to screen and identify input sequences as therapeutic from non-therapeutic proteins/peptides. In this module users can submit both single and multiple protein sequences in FASTA format. Secondly, the “motif scan module” may be used to separate therapeutic proteins/peptides from non-therapeutic proteins/peptides based on exclusive motifs identified using the MERCI tool. It scans for patterns in the user-provided query sequence. Further, the ‘therapeutic protein design module’ helps the user by converting the user query sequence to therapeutic proteins/peptides with possible single point mutation in their input sequence. This web server is developed with a responsive HTML template and compatible with several operating systems. In addition to this, we also provide a ThpPred standalone Python package, which can be accessed using the web server’s “Download” section, to let users forecast therapeutic proteins/peptides at a larger scale.

## Discussion

In the last two decades, the rapid advancement in next generation sequencing technologies has resulted in accumulation of nearly 250 million protein sequences in the UniProt repository [https://www.ebi.ac.uk/uniprot/TrEMBLstats]. Screening of therapeutic potential of proteins/peptides available at public repositories is the need of the hour, for identifying novel drug candidates. In order to facilitate this, ThpPred is a novel method developed for predicting therapeutic potential of a given protein/ peptide sequence. In this study, we have made a systemic approach and utilized FDA approved therapeutic proteins as a positive dataset, to have higher accuracy in our predictions. We have incorporated standard protocols that are widely accepted in literature as best practices in our tool. In addition to this, a web-based platform and standalone package is provided to enable users to categorize therapeutic and non-therapeutic proteins/peptides as per their sequence data. We believe that our study will be useful for the scientific community working in designing protein-based therapeutics and drugs. This tool will enable them to do virtual screening of potential candidates and would save time and cost involved in the field of protein/ peptide-based therapeutics. It is envisaged that the researchers will make considerable use of this prediction approach to develop more effective and precise protein-based therapies for treating a range of ailments.

## Future Scope

In the current study, we have implemented machine learning models and motif information for accurately predicting therapeutic and non-therapeutic proteins/peptides. In future we plan to club physicochemical properties of therapeutic proteins/peptides along with the machine learning models to enhance our predictability.

## Author Contributions

SJ and SG collected and processed the datasets. SJ, SG and GPSR implemented the algorithms. SJ and SG developed the prediction models. SJ and GPSR analyzed the results. SP and SG created the web server interface. SJ, SG and GPSR penned the manuscript. GPSR conceived and coordinated the project, and gave overall supervision to the project. All authors have read and approved the final manuscript.

## Acknowledgement

SJ and SG are thankful to the Department of Computational Biology, IIIT-Delhi for infrastructure and facilities.

## Data Availability Statement

All the datasets generated for this study are available at the “ThPPred” web server, https://webs.iiitd.edu.in/raghava/thppred/download.php. The source code is hosted on GitHub and can be found at https://github.com/raghavagps/thppred.

